# Cell-cell contact dictates life or death decisions following CD95 activation in cancer

**DOI:** 10.1101/308346

**Authors:** Gülce S. Gülcüler Balta, Cornelia Monzel, Susanne Kleber, Joel Beaudouin, Thomas Kaindl, Meinolf Thiemann, Christian R. Wirtz, Motomu Tanaka, Ana Martin-Villalba

**Affiliations:** Department of Molecular Neurobiology, German Cancer Research Center (DFKZ), Im Neuenheimer Feld 581, 69120 Heidelberg, Germany; Faculty of Biosciences, Heidelberg University, 69120 Heidelberg, Germany; Physical Chemistry of Biosystems, Institute of Physical Chemistry, Heidelberg University, 69120 Heidelberg, Germany; Experimental Medical Physics, Heinrich-Heine University Düsseldorf, 40225 Düsseldorf, Germany; Department for Bioinformatics and Functional Genomics, Bioquant and Institute of Pharmacy and Molecular Biotechnology, Heidelberg University, 69120 Heidelberg, Germany; Division of Theoretical Bioinformatics, German Cancer Research Center (DKFZ), Im Neuenheimer Feld 267,69120 Heidelberg, Germany; Apogenix AG, Im Neuenheimer Feld 584, 69120 Heidelberg, Germany; Department of Neurosurgery, University Hospital Ulm, 89312 Günzburg, Germany; Center for Integrative Medicine and Physics, Institute for Advanced Study, Kyoto University, 606-8501 Kyoto, Japan

## Abstract

Cancer cells react to CD95 activation with either apoptotic or tumorigenic responses. Yet, the determinants of these two antithetic reactions are fundamentally not understood. Here, we show that pre-confined CD95L molecules activate apoptosis of cancer cells *in-vitro*. For particular CD95L pre-confinement, apoptosis activation is most efficient. Surprisingly, in tumor models, the same pre-confinement yields enhanced proliferation of cancer cells. This shift is rooted in cell-cell interactions, as proliferation was also observed in tumorspheres *in-vitro*. Indeed, proliferation required death-domain tyrosine phosphorylation of CD95 that was facilitated by cell-cell contacts, whereas decreasing the levels of global tyrosine kinase activity favored apoptosis. Altogether, the response to CD95 activation is cell context-dependent and tunable by CD95L pre-confinement, thereby opening therapeutic opportunities in cancer.

**One Sentence Summary:** Cell-cell contact tunes tyrosine-kinase activity thereby dictating life or death upon CD95 activation by pre-confined CD95L.

## Introduction

CD95 (Fas/ APO-1/ TNFRSF6) belongs to the tumor necrosis factor (TNF) receptor subfamily of death receptors, which is characterized by a cytoplasmic sequence termed the death domain (DD). Discovered as the prototypic trigger of the apoptotic pathway upon binding to its cognate ligand, the CD95-ligand (CD95L) (1–4), follow-up studies revealed that most primary tumor cells were resistant to apoptosis (5, 6). The activation of apoptosis by CD95 requires the assembly of a death inducing signaling complex (DISC) composed of the adaptor protein Fas-associated protein with death domain (FADD), and the initiator caspases-8 and −10. DISC assembly further activates other effector caspases in charge of cleaving cell-substrates leading to the cell’s own demise (7).

Surprisingly, the CD95-mediated increase in tyrosine kinase activity reported in the first studies was not followed up (8). It was only in the last decade that several studies reported a myriad of cell responses ranging from migration, survival, differentiation, to proliferation downstream of CD95 activation (9, 10). Accordingly, other signaling pathways such as activation of NF-κB and ERK were linked to CD95-mediated motility and invasiveness of cancer cells (11). We previously showed that a tyrosine residue within a immuno receptor tyrosine-based inhibition/activation-like motif (ITAM/ITIM) in the DD of CD95 is susceptible to phosphorylation by Src family kinases (SFK), thereby serving as a protein-protein interaction platform involved in the recruitment of SH2-domain containing proteins that lead to downstream kinase activities such as ERK and PI3K (6, 12, 13). This differential CD95 activation mode has been linked to different forms of CD95L, the membrane-anchored (mCD95L) or soluble form (sCD95L) (14–17).

## Results

To study the activation of CD95 receptor by mCD95L, we designed membrane-based surrogate cell surfaces (18, 19) displaying the soluble CD95L fused to the T4-Foldon motif used as a trimerization domain (Fig. 1A) (12). This membrane model system enables quantitative and precise control over local concentrations of CD95 signaling components, since the average lateral CD95L density is tuned by changing the doping ratio of biotinylated lipids in the lipid membrane (Table S1).

**Figure 1.**
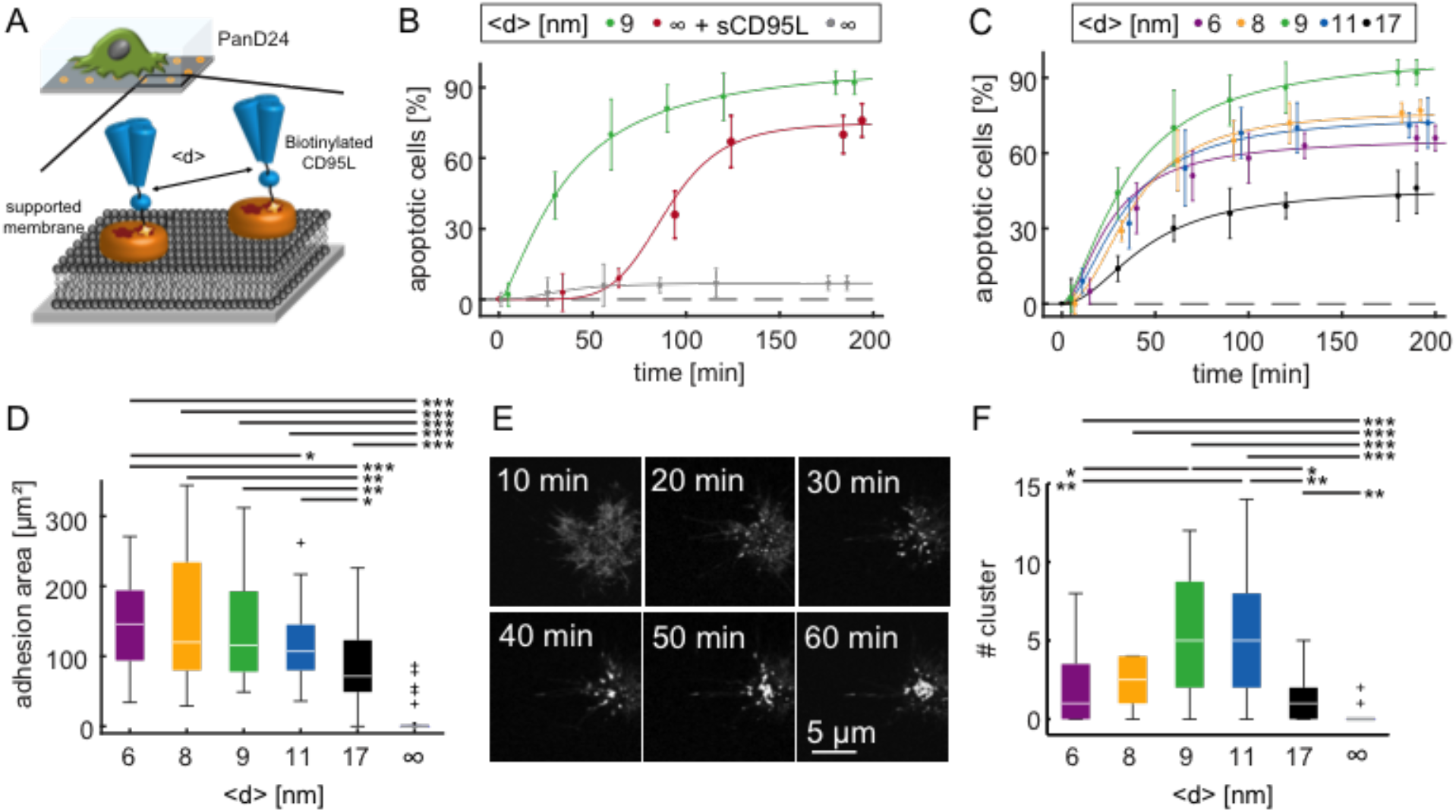
Optimal induction of apoptosis in PanD24 cells occurs at specific CD95L intermolecular distance and correlates with CD95 cluster formation. (A) Scheme of the experimental setup with PanD24 cells on supported membranes displaying CD95L at defined average intermolecular spacing <d>. n ≥ 90 cells/condition. (B) Comparison of apoptosis induction at optimal ⟨*d*\*⟩ = 9 nm, in the presence of soluble CD95L, and in the absence of CD95L in PanD24 cells. (C) Dynamics of apoptosis induction in PanD24 cells at different <d>. The percentage of apoptotic cells over time is analyzed via a Hill fit (Eq.1, Table S2). (D) Cell adhesion area vs. <d>, recorded at t = 2 h. (E) Fluorescence images of PanD24 membrane contact area of CD95-YFP transfected PanD24 cells at <d> = 9 nm. Significance levels according to p***< 0.001, p**< 0.01, p*< 0.05. (F) Number of CD95 clusters (≥ 1 μm^2^) vs. <d>, recorded 10 min before apoptotic blebbing.

To test the function of this mCD95L, we exposed cancer stem cells isolated from a pancreatic ductal adenocarcinoma (PDAC) patient (PanD24) to the CD95L-membrane and monitored their apoptotic response over time using label-free time-lapse microscopy. The average intermolecular distance between ligands is defined as ⟨*d*⟩ in the rest of the text. Control membranes without CD95L are defined as ⟨*d*⟩=∞. Using a ligand density of 13200 ligands per μm^2^, corresponding to ⟨*d*⟩ = 9 nm, we observed a rapid rise of the number of apoptotic cells, which reached 92% with a half time of 36 min (Fig. 1B). In contrast, 7% of apoptotic cells were observed on membrane with no ligand, confirming that apoptosis results from specific CD95-CD95L interactions. Moreover, the same ligand used in solution at 1 μg/ml also lead to apoptosis reaching 70% but with slower kinetics and showing a lag time of 40 minutes before an abrupt rise of the number of apoptotic cells. The number of apoptotic cells over time followed a sigmoid curve, which was fitted with a Hill-equation (solid lines) to characterize the dynamics of the cellular response:

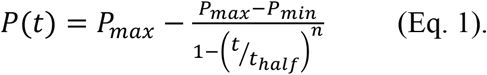

*P_min_* and *P_max_* are the minimum and maximum percentages of apoptotic cells, ⟨*t_halif_*⟩ is the characteristic time for apoptosis induction after CD95L contact, and *n* is the Hill coefficient which indicates how efficiently cells become apoptotic, and by extension how the extracellular stimulus is recognized. The fit yielded a Hill coefficient of *n* > 3 for the soluble ligand compared to *n* ∼ 1.5 for mCD95L. This indicates that the apoptotic cascade induced by sCD95L requires a long process of signal integration that is strongly reduced by the membrane form of the ligand.

This observation leads to the hypothesis that the topology and density of the membrane ligand can activate the receptor extremely efficiently and trigger rapid apoptosis. To further test this hypothesis, we measured the apoptotic response for different ligand densities, with an average ⟨*d*⟩ ranging from 6 to 17 nm. All tested densities led to a rapid increase of the number of apoptotic cells over time (Fig. 1C). Strikingly, the number of dead cells and the apoptosis efficiency did not increase monotonically with the density of ligand but showed an optimum towards 9 nm (Fig. 1C and Table S2).

This optimal ligand distance likely corresponds to the most favorable one for the ligand-receptor interaction, and we therefore further investigated this interaction. First, by measuring the cellular surface on the membrane, we observed that the optimal ligand distance does not correlate with a maximal cell adhesion (Fig. 1D). On the contrary, this adhesion increases monotonically with the ligand density. Second, we observed the organization of the receptor on the plasma membrane by expressing CD95-YFP in the PanD24 cells and imaging their fluorescence at the level of the CD95L-membrane (Fig. 1E). While at early time points, the cell spreading is accompanied by a diffuse YFP signal (Fig. 1E, 10 min), bright clusters of receptor appeared and accumulated over time to areas exceeding 1 μm^2^ (Movie S1). Strikingly, the number of clusters, measured 10 min before the cells died, was maximum for <d> = 9 to 11nm, correlating well with the optimal efficiency of apoptosis (Fig. 1F). Thus, the optimal distance (defined as ⟨*d*\*⟩) of ligand likely leads to ligand-receptor interactions and cluster formation that induce particularly strong apoptotic signaling. The ⟨*d*\*⟩ = 9 to 11 nm relates more to the scale of proteins than to a typical density of a given receptor on the plasma membrane. In PanD24, we estimated the density of CD95 receptors to 13.4 per μm^2^ (Table S3), 1000 times less than that of the ligand on the supported membrane. So, we hypothesized that the optimal ligand distance correlates with a critical size of the receptors and should therefore be universal and independent of the receptor density. To address this point, we examined primary cancer cells originating from different GBM tumors (20). GBM60 and GBM50 cells were isolated from two different patients and both expressed 1.8 receptor per μm^2^, lower number than PanD24 (Table S3). With a ligand intermolecular distance of 11 nm, both cell lines responded with a kinetic of apoptosis that was similar to the one of PanD24 upon CD95L exposure (Fig. 2A). However the final number of apoptotic cells was lower, with a plateau at around 35% for GBM60 and 21% for GBM50, while PanD24 reached 75%. We repeated the characterization of the death curves at different ligand densities for the two cell lines and we observed that, like for PanD24, the distance ⟨*d*\*⟩ = 9 to 11 nm lead to the maximum number of apoptotic cells, the fastest kinetics of apoptosis (Fig. 2B) and the smallest Hill coefficient (Table S4). Taken together, these data suggest the existence of a universal optimal CD95L spacing at 9 to 11 nm to trigger apoptosis most efficiently.

**Figure 2.**
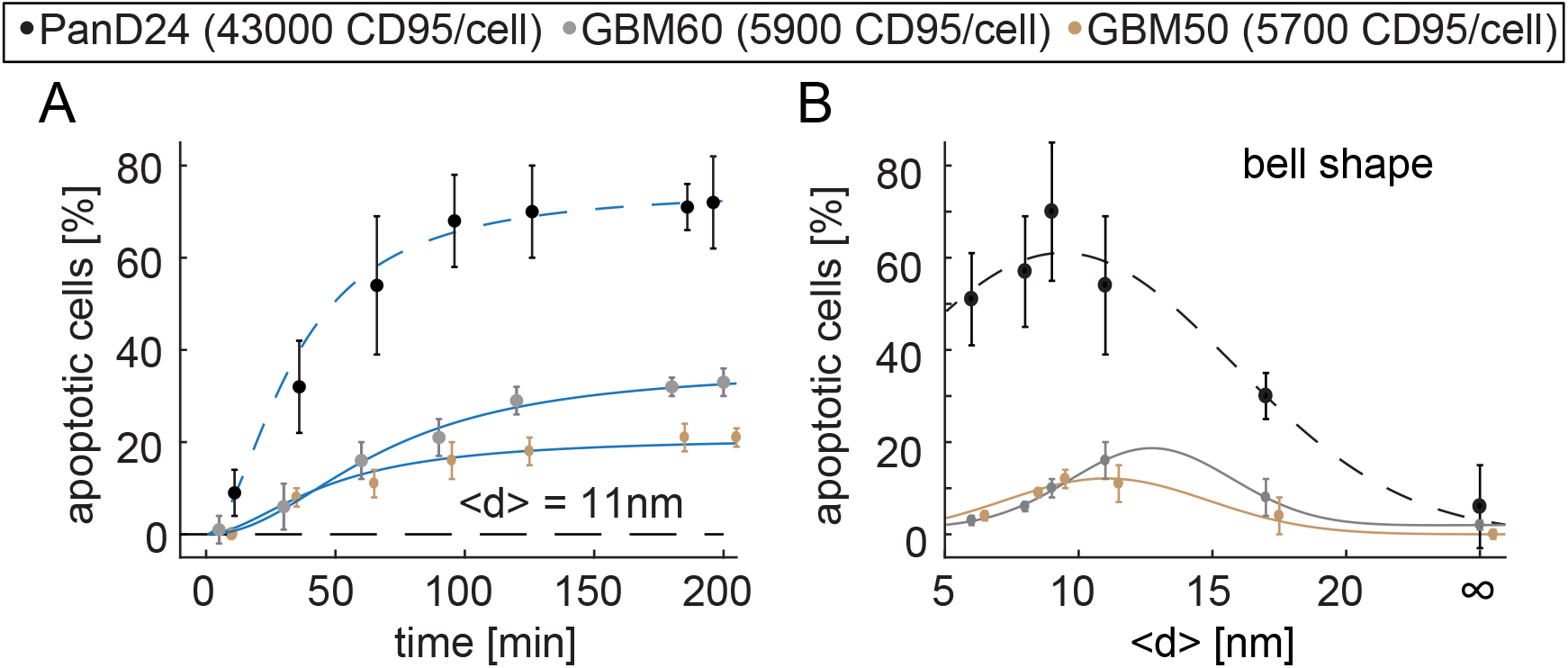
Apoptosis of primary cancer cells correlates with CD95 expression levels and exhibits an optimal CD95L distribution as a generic signature. (A) Dynamics of GBM apoptosis at <d> = 11 nm. n ≥ 70 cells/condition. (B) Percentage of apoptotic cells vs. <d> (t = 1 h). Data of PanD24 at <d> = 11 nm are indicated as broken lines as reference.

Furthermore, to decipher whether the difference in signaling dynamics and efficiency stems from the mobility of CD95-CD95L pairs and their re-arrangement on the substrate, we prepared membranes facilitating or suppressing the lateral diffusion of ligands. While the experiments described so far were performed with DOPC based membranes, where CD95L-CD95 pairs can diffuse, we now used DPPC based membranes, where lateral lipid diffusion is significantly suppressed. This arises from the DPPC lipid chain melting transition temperature at 41 °C, which, at the experimental temperature of 37°C, corresponds to the lipid gel phase. In the latter, lateral diffusion coefficients are 2-3 orders of magnitude smaller than those measured in the liquid-crystalline phase (21). On these DPPC membranes, GBM50 and GBM60 cells died much slower than on DOPC (Fig. 3B, F versus Fig. 3A, E). However, the maximum level of apoptotic cells after 6 h reached the same values in GBM cells exposed to DPPC membranes as compared to those obtained on DOPC membranes after 3 h. The distinct decrease in the fraction of apoptotic cells was further accompanied by a larger Hill coefficient on DPPC (*n* = 3 – 6, Table S5) with respect to DOPC membranes (*n* = 1.7 – 4, Table S4). Accordingly, a suppression of CD95-cluster formation (Fig. 3D, H and Fig. 3C, G) was observed, indicating that loss of lateral diffusivity suppressed the formation of death-inducing clusters. Notably, the most efficient apoptosis and lowest Hill coefficients were again recovered for the optimal spacing of 9 to 11 nm for both cell and membrane conditions.

**Figure 3.**
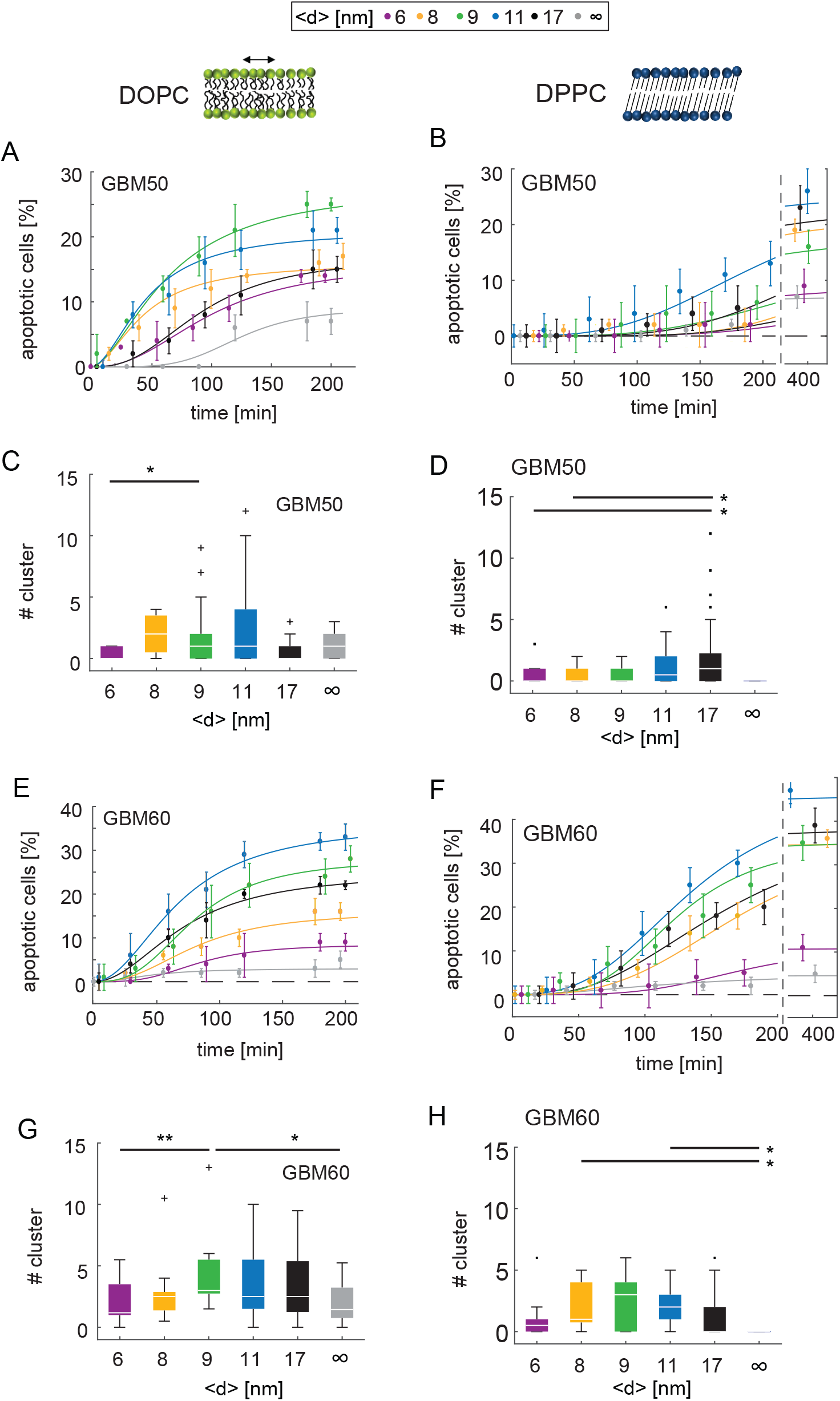
Apoptosis induction in GBM cells is optimal at specific CD95L intermolecular distances on DOPC supported membranes and correlates with CD95 cluster formation. Inhibition of CD95L mobility via DPPC supported membranes results in slowdown of apoptosis signaling and suppression of CD95 cluster formation. (A,E) Dynamics of apoptosis induction in GBM50 and GBM60 cells at different <d> on DOPC. The percentage of apoptotic cells over time is fitted with a Hill function (Eq.1, see Table S4 and text for fit results). (C,G) Number of CD95 clusters on DOPC supported membranes at different <d>. Clusters are ≥ 1 μm^2^ and their number is analyzed 10 min before cell blebbing/apoptosis. n≥ 40 cells/condition. (B,F) Dynamics of apoptosis induction in GBM50 and GBM60 cells at different <d> on DPPC (see Table S5 and text for fit results). (D,H) Number of CD95 clusters on DPPC supported membranes at different <d>. Significance levels according to p*** < 0.001, p** < 0.01, p* < 0.05.

So far, our data show that CD95L placed at an optimal intermolecular distance can successfully trigger CD95 mediated apoptosis in cancer cells *in vitro*. Thus, we hypothesized that this knowledge could be used to effectively kill tumor cells *in vivo*. To this end, we assembled beads coated with supported membranes and decorated with CD95L at the optimal intermolecular distance of 9 nm, as well as without CD95L (Fig. 4A). In addition, patient-derived GBM cells were FACS sorted according to their CD95 surface expression levels, as previously described (20). CD95^high^, CD95^low^ or unsorted tumor cells (bulk) were then subcutaneously (s.c.) injected into the flanks of immunosuppressed SCID^bg^ mice (Fig. 4A). S.c. location was selected over the brain to avoid beads having to cross the blood brain barrier to reach the tumor. Three weeks after cell injection, beads were intravenously (i.v.) injected and tumor growth was further monitored. First, the beads spread systemically, as they were found in several locations such as the tumor, lung and liver (Fig. S1). Notably, there were no signs of apoptosis in lung and liver but rather induction of inflammatory infiltrates, as previously reports (22). To our surprise, treatment with CD95L-beads exponentially increased growth of bulk- or CD95^high^-tumors but not of CD95^low^-tumors or tumors treated with control-beads (Fig.4B). Additionally, the same CD95-dependent growth behavior was observed in xenograft tumors derived from another GBM patient (Fig. S2). This finding demonstrates that beads coated with CD95L at the optimal distance foster proliferation of tumor cells *in vivo*.

**Figure 4.**
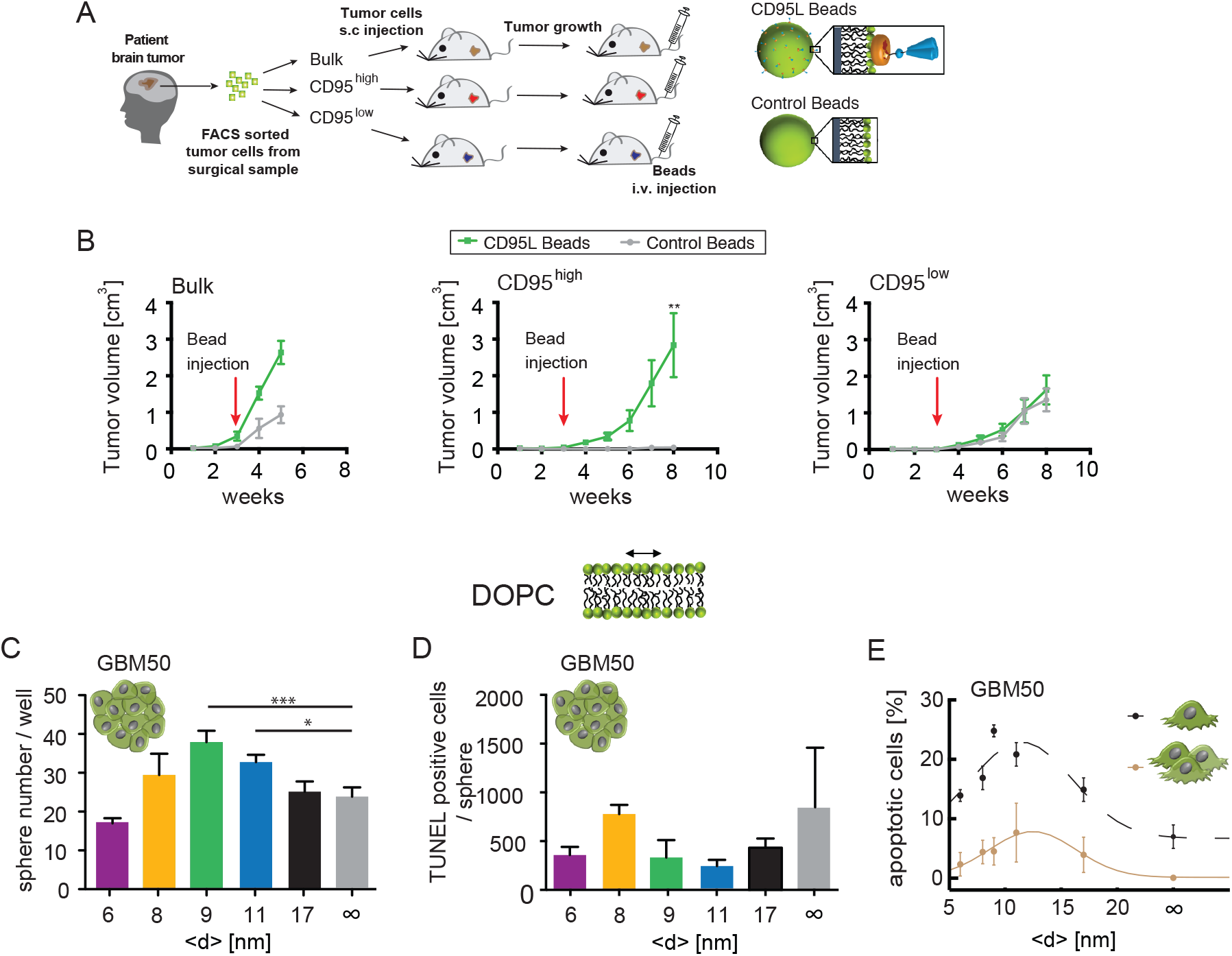
CD95L coated beads promote subcutaneous tumor growth and self-renewal of tumor-spheres. (A) Scheme of the patient-derived tumor xenograft model. Cancer cells are FACS sorted according to CD95 surface expression levels and grown in mice subcutaneously before the beads i.v. injection. Either non-functionalized DOPC membrane coated beads (control beads) or CD95L functionalized beads were injected. (B) Kinetics of tumor progression upon bead injection (control beads = grey data, n = 3; CD95L beads = green data, n = 4 – 5). CD95L distance on beads <d> = 9 nm. Values are mean ± SEM; two-way ANOVA p** < 0.0013. (C) Patient derived glioma sphere cultures (GBM50) were treated for 4 h with CD95L-coated membranes and were subsequently plated for sphere formation assessment. Values are mean ± SEM.; one-way ANOVA p* < 0.02, p*** < 0.0002. (D) TUNEL assay for patient derived glioma sphere cultures (GBM50) was performed to assess apoptotic cells after 4h treatment with CD95L-coated membranes. Values are mean ± SEM. (E) Comparison of apoptotic cell percentage in single cells vs cells in contact over different intermolecular ligand distances.

What is the origin of these antithetic cellular responses observed *in vivo* and *in vitro*? One apparent difference is the cell context *in vivo* given by the multicellular network of the tumor vs. isolated cells on planar surfaces *in vitro*. To test if the different cell behaviors originate from cell-cell interactions, we investigated the behavior of cancer stem cells within tumorspheres *in vitro*. Self-renewal was assessed by counting the number of tumorspheres as a function of ⟨*d*⟩. Surprisingly, also *in vitro*, exposure of tumorspheres to CD95L-supported membranes but not control-membranes (⟨*d*⟩=∞) triggered a significant increase of self-renewal (Fig. 4C) rather than apoptosis (Fig. 4D). Importantly, the maximum sphere number (38 ± 3) was found at the optimal distance ⟨*d*\*⟩ (Fig. 4C). This behavior was also observed in PanD24 tumorspheres (Fig. S3). Based on our data, we conclude that the confinement of CD95L at ⟨*d*\*⟩ enhanced the self-renewal most efficiently in both solid tumors *in vivo* and tumorspheres *in vitro*. We then addressed if the presence of mere one or two cell-cell contacts would be enough to shift the response of the cell to CD95 activation away from apoptosis. Therefore, we monitored and compared apoptosis of GBM50 cells either exhibiting up to two cell-cell contacts or as isolated cells on the same supported membranes (Fig. 4E, Movie S2). GBM cells in contact with other cells exhibited a distinctly lower fraction of apoptosis induction (reduction by ∼ 2 fold) for all ⟨*d*⟩ conditions as compared to isolated cells. Moreover, the cells from other GBM patients, as well as Hela cells from another cancer type (cervical cancer) showed lower apoptosis when they engaged to the ligand as a group of cells (Fig. S4, broken lines). In summary, in solid tissues where cells are in contact with neighboring cells, CD95 activation promotes pro-survival signaling, whereas loss of cell-cell contact renders CD95-stimulated cells prone to undergo apoptosis.

To shed light on the cross talk between the signaling at cell-substrate and cell-cell contacts, we performed the experiments on supported membranes composed of DPPC. The apoptotic behavior of single vs. cells in contact was examined on DOPC or DPPC substrates (Fig. S5). GBM cells in contact reached similar maximal levels of apoptotic rate in DPPC and DOPC membranes though with a narrower range of optimal distance on DPPC substrates. These data demonstrate that the inhibition of apoptosis by cell-cell contacts is not influenced by the mobility of CD95L.

To gain further mechanistic insights into pro-tumorigenic signaling of CD95 in the natural environment of the brain with a fully competent immune system, we used a syngeneic orthotopic glioma model (Fig. 5A). To this end, cells derived from a spontaneous murine astrocytoma (SMA560) were intracranially (i.c.) injected into the striatum of VMDK mice. Five days post tumor cell implantation, control beads (coated with DOPC, no CD95L) or CD95L beads (coated with biotin lipid doped DOPC membranes displaying CD95L at ⟨*d*⟩ = 8 nm) were i.c. injected. The mice were sacrificed once the maximum tolerable tumor volume was reached. Thereafter, brain tumor sections were stained for phosphorylated ERK (p-ERK), phospho Histone3 (p-H3), cleaved Caspase-3, and phosphorylated Akt (p-Akt) (Fig. 5B). The number of cells positive for p-ERK and p-Akt were significantly higher (Fig. 5C) in tumors exposed to CD95L beads, than control bead-treated tumors. This is in line with previous studies, demonstrating that the pro-tumorigenic CD95-signaling cascade involves activation of the ERK and/or PI3K signaling pathways (9, 12, 23). A clear increase in the mitotic index of tumors treated with CD95L beads compare to control was assessed by p-H3 staining (Fig. 5B-C). On the other hand, there was no difference in the number of cells undergoing apoptosis between the two groups, as assessed by staining of cleaved Caspase-3 (Fig. 5B-C). Hence, CD95-dependent proliferation of cancer cells highly correlates with increased activation of Ras/ERK and PI3K signaling pathways.

**Figure 5.**
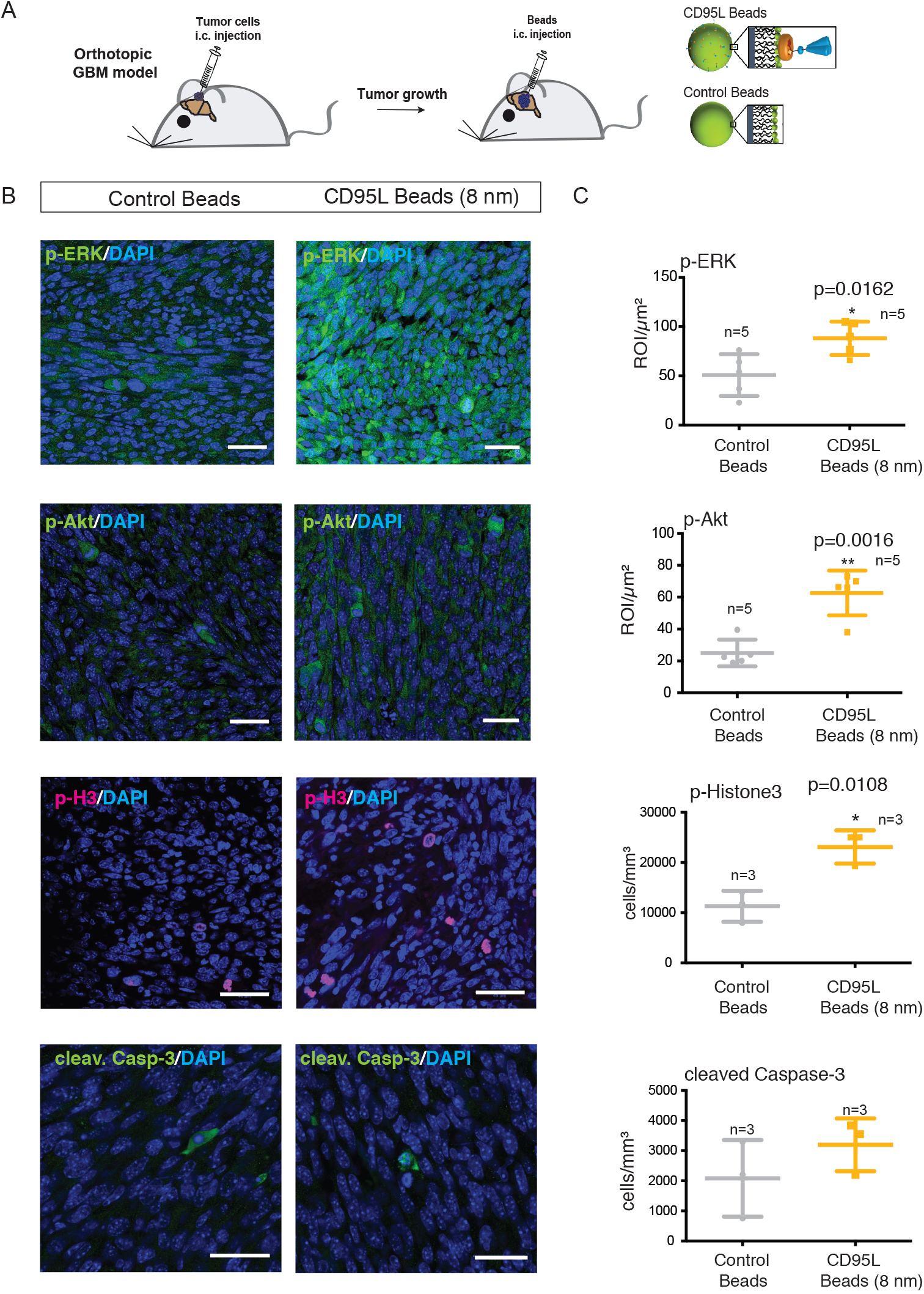
CD95 activation induces phosphorylation of ERK, and enhances mitosis as non-apoptotic functions in syngenic orthotopic glioma tumors. (A) Syngenic tumor model. (B) Representative confocal images for p-ERK (n = 5) and p-AKT (n=5), p-Histone3 (n = 3) and cleaved Caspase-3 (n = 3), scale bar 30 μm. Left; control beads, right; CD95L-functionalized beads (<d> = 9 nm). (C) Quantification of antibody stained tumor sections. Cell density (cells/mm^3^) calculated from confocal images. For quantification of p-ERK and p-Akt signal intensity (ROI) per confocal image was plotted. Values are mean ± S.D.; t-test p*<0.05, p**<0.01.

We have previously reported that CD95 activates these pathways through recruitment of SH2-adaptor proteins to a phosphorylated tyrosine residue within its DD (9). Thus, signaling modules involved in death or proliferation compete at the level of the DD, as previously shown in pancreatic adenocarcinoma cells (23). To test the importance of this tyrosine motif in CD95-mediated cellular responses, we generated GBM60 cells stably overexpressing either CD95-alanine mutant receptors (CD95-Ala) that substitute the Y291 residue of the human CD95 with alanine, or wildtype CD95 (CD95-wt). Upon exposure of these cells to CD95L-surrogate membranes, cells overexpressing the CD95-Ala showed a higher induction of apoptosis compared to the cells overexpressing CD95-wt (Fig. 6A). Moreover, the induction of sphere formation upon CD95L membrane stimulation was blocked by CD95-Ala overexpression (Fig. 6B). This data highlights the critical regulation of apoptosis and survival at the DD of CD95 and the requirement of tyrosine kinase activity to bias CD95 cell responses toward cell proliferation. Hence, we hypothesized that cell-cell contact might modulate the levels of tyrosine-kinase activity. Indeed, the presence of cell-cell contact globally increased levels of tyrosine-kinase activity as compared to levels exhibited by single cells (Fig. 6C).

**Figure 6.**
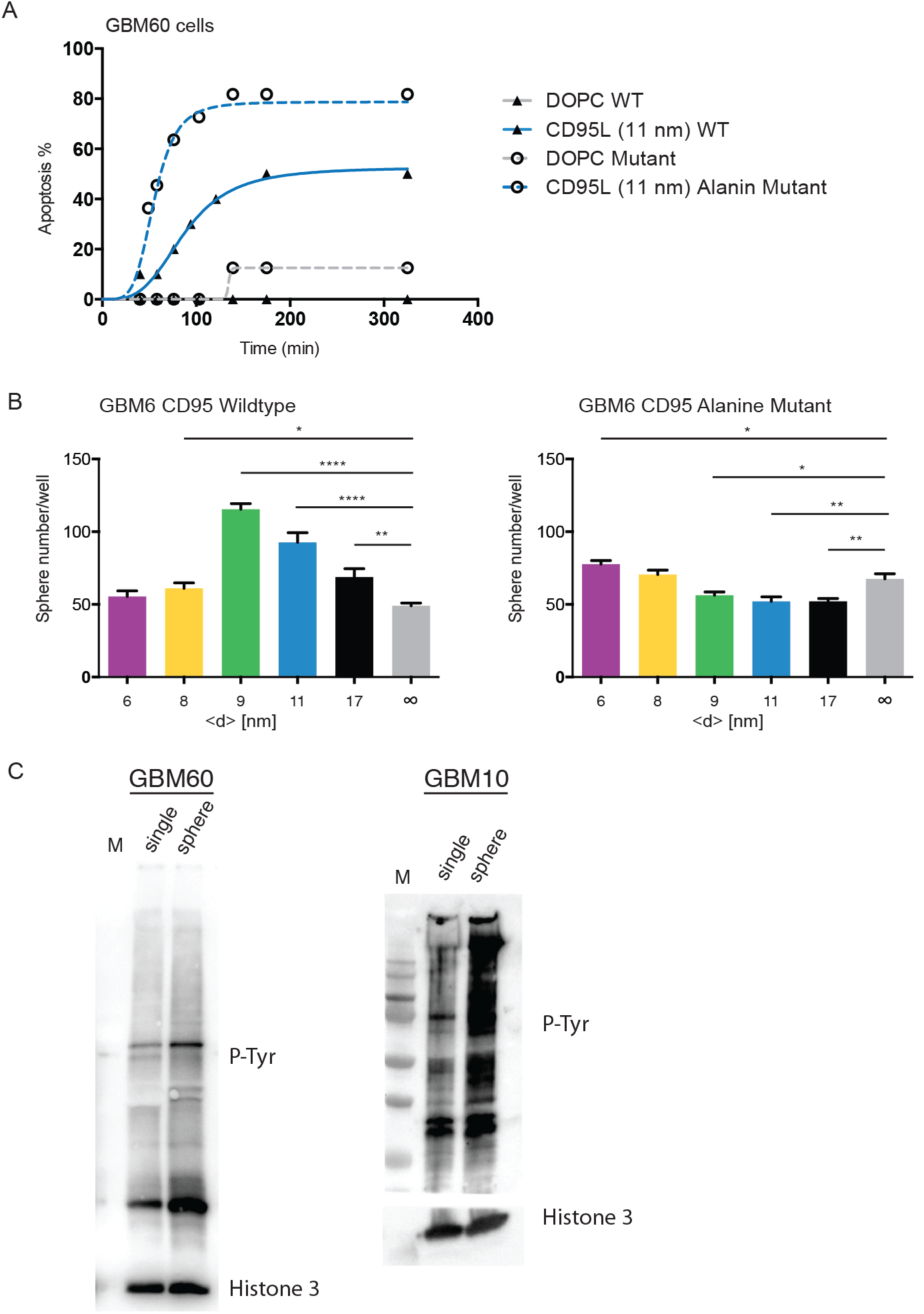
CD95 activation induces phosphorylation of ERK and thereby enhances proliferation. (A) Quantification of apoptosis induction in CD95-Ala mutant compared to CD95-wt cells by CD95L membranes. The percentage of apoptotic cells over time is analyzed by fitting a Hill function. (B) Glioma sphere cultures (GBM6) over-expressing CD95-wt or CD95-Ala mutant were treated for 4h with CD95L-coated membranes at different <d>. Subsequently they were plated to measure the sphere forming capacity. Values are mean ± SEM; t-test p* < 0.05; p**** < 0.0001. Western Blot analysis of single cells versus spheres from GBM60 and GBM10, probed for pTyr antibody. Anti-Histone antibody was performed on the same membranes and used as a loading control.

## Discussion

In this study, we have established membrane-based surrogate cell surfaces displaying CD95L to study CD95 activation in cancer cells. This system has unveiled an optimal CD95L intermolecular distance as efficient trigger of CD95 cluster formation with subsequent apoptosis in primary cancer cells. In particular, pre-confinement of CD95L on the membrane at an intermolecular distance ⟨*d*\*⟩ = 9 – 11 nm was found to maximize the efficiency and kinetics of receptor activation. The evolution of CD95 clusters was found to scale with apoptosis induction in a similar manner. Recent PALM imaging confirmed the existence of CD95 clusters and showed their dependence on palmitoylation of CD95 for lipid-raft localization (24). A rigid-body based computational model of membrane receptor-ligand interactions further illustrated that confining the structural flexibility of receptors via membrane topology has an influence on the binding to its ligand (25). Modification of intramolecular flexibility could optimize the binding between ligand-receptors. Indeed, our model system not only demonstrates the relevance of the intermolecular spacing of ligands, but also shows that ligand orientation through membrane tethering has an essential influence on the cellular response in comparison to soluble ligands. Particularly surprising was our discovery that cell-cell contact can switch CD95 activation from apoptosis to tumorigenesis. Along this line, increased cell-cell contact via control of cell spreading using micropatterned substrates increased proliferation of muscle and endothelial cells (26). More recently, in a cell intercalation study in the *Drosophila pupa*, the weakening/loosing of cell contacts was shown to result in reduction of PIP3 levels, which in turn stimulated the loser phenotype, and elimination of these cells by apoptosis (27). Additionally, cell-cell contact might also have a direct influence on membrane receptors. Cell-cell contacts promoted vesicular recycling of EGFR and maintained ERK activity in cancer cells leading to switch from migration to proliferation (28). On a molecular scale, cell-cell contact may energetically favor segregation of proteins or result in the formation of multiprotein complexes, thus resulting in different molecular stoichiometry, affinity of binding, and energetic losses from membrane bending (29–31). Accordingly, we tested how phosphatase/kinase activity is modulated in our experiment. We showed that cell-cell contact increases tyrosine-kinase activity. The basis of the global tyrosine-kinase activity could be the transactivation of CD95 which was reported for Integrins (32) and RTKs like EGFR (33).

Altogether, this study indicates that the switch from apoptosis to survival is not determined by the CD95-CD95L interaction, but rather by the cell context. Blocking non-apoptotic functions of CD95 would be a promising therapy option for cancer. Accordingly, a Phase II clinical trial of a fusion CD95-FC protein was shown to neutralize CD95 activity in glioblastoma patients, which increased the overall survival of patients with unmethylated-CD95L promoter (34). Yet, as reported for previous strategies targeting kinase activities, downstream of growth receptors, inhibition of CD95 might be compensated by other tyrosine-kinase receptors in the long term (35). Our study unveils a new strategy to convert the tumorigenic signal into apoptosis by targeting tyrosine phosphorylation at DD of CD95 and thereby avoids the risk of therapy resistance in cancer patients.

## Acknowledgments

We thank the Nikon Imaging Center at the University of Heidelberg for providing assistance in their imaging facility. We thank Katrin Volk for technical support. Histology services for tumor pathology analysis were provided by the Institute of Pathology, University Hospital Heidelberg.

## Author contributions

Conceptualization; A.M.-V., M.T., Investigation; G.S.G.B., C M., S.K., J.B., T.K., Resources; Me.Th., C.R.W., Writing – Original Draft Preparation; G.S.G.B., A.M.-V., M.T., Writing – Review & Editing; C.M., J.B., S.K., Supervision; A.M.-V., M.T.

## Data and materials availability

All data is available in the main text or the supplementary materials.

## Materials and Methods

### Materials

1,2-dioleoyl-*sn*-glycero-3-phosphocholine (DOPC), 1,2-dipalmitoyl-*sn*-glycero-3-phosphocholine (DPPC), and 1,2-dioleoyl-*sn*-glycero-phospho-ethanolamine-3-N-(capbiotinyl) (biotin-DOPE) were purchased from Avanti Polar Lipids (Alabaster, AL, USA). Neutravidin (deglycosylated avidin) was purchased from Life Technologies (Carlsbad, CA, USA). Biotinylated CD95L trimer (CD95L-T4-btn) was custom synthesized by Apogenix AG (Heidelberg, Germany). Media and cell culture supplements were bought from Life Technologies and all other chemicals in p.A. quality from Sigma-Aldrich (St. Louis, MO, USA), unless specified otherwise. Throughout this study, de-ionized ultrapure water (Genpure, TKA Niederelbert, Germany) was used for preparation of all buffers and vesicle solutions.

### Culture of PanD24 Cells

Pancreatic cancer cells (PanD24) were isolated from a male patient suffering from an invasive malignant pancreas tumor. Collection of patient samples was approved by the Ethics committee of Heidelberg University, Medical Faculty (S-604/2012). Cells were cultured in sterile filtered DMEM:Ham’s nutrient mixture F12 media (1:1; Thermofisher, Waltham, MA, USA) supplemented with 2 mM L-glutamine, 1 vol% penicillin and streptomycin (PenStrep; Thermofisher), 10 vol% N2 medium supplements, 0.5 vol% B 27 medium supplements, 20 ng/ml basic human fibroblast growth factor (bFGF), and 20 ng/ml epidermal growth factor (EGF). Cells grew non-adherently and were split by a factor of 1:8 as soon as agglomerates were visible by eye. Prior to experiments, cells were washed twice in 150 mM phosphate buffered saline (PBS) at pH 7.4 and centrifuged. Experiments were performed at concentrations of ∼ 2 × 10^5^ cells/ml as single cell suspension in freshly prepared culture medium without phenol red and PenStrep.

### Culture of patient-derived glioblastoma (GBM) cells

Collection of patient samples was approved by the Ethics committees of the Charité University Medicine, Campus Virchow-Klinikum, Berlin (EA3/023/06) and Heidelberg University, Medical Faculty (S-233/2006). The specimen was examined by a neuropathologist to confirm that the tumor met WHO criteria for glioblastoma. The sample was dissociated using the Brain Tumor Dissociation Kit (P) (Miltenyi Biotec GmbH, Bergisch-Gladbach, Germany) and subsequently expanded in Neurobasal medium supplemented with B27 (2 vol.%), 2 mM L-glutamine, heparin (2 μg/ml), EGF (20 ng/ml) and bFGF (20 ng/ml) at 5% CO_2_ and 37°C.

Glioblastoma multiforme (GBM) stem cells were established from mechanically and enzymatically dissociated brain tumor samples. The cells were cultured in serum-free medium containing stem-cell mitogens as mentioned above. Prior to experiments, the agglomerated cells were centrifuged and re-suspended with Accutase (Thermofisher) to get a single cell suspension. Then, the cell suspension was diluted in freshly prepared cell culture medium to concentrations of ∼ 2 × 10^5^ cells/ml.

### Culture of SMA-560 and Hela cells

SMA560 and Hela cells were maintained in DMEM supplemented with 10% fetal calf serum, 1x glutamine and 1x penicillin/streptomycin. Cells were kept in 5% CO_2_ at 37°C.

### Lentiviral infections

For PanD24, cells were infected with lentiviral vector pEIGW-CD95-YFP at a multiplicity of infection (MOI) of 5. Expression of all transgenes was confirmed in infected cells by FACS analysis of YFP expression.

For lentiviral infections, GBM cells were sorted for CD95 negative surface expression (CD95^neg^ cells). CD95^neg^ cells were infected with the lentiviral vector pEIGW-CD95-Tyr (Y291) or pEIGW-CD95-Ala (A291) at a multiplicity of infection (MOI) of The plasmids were constructed by site directed mutagenesis to exchange the tyrosine residue (Y291) to alanine in the DD of CD95 in pEIGW-CD95-Tyr. All lentiviruses were propagated using previously described methods (36). Expression of all transgenes was confirmed in infected cells by FACS analysis of GFP expression. The percentage of infected cells was 60-80%. The GFP positive cells were sorted according to their expression level and expanded for further experiments.

### Preparation of supported membranes

Before the assembly of sample chambers, glass substrates were cleaned by a modified RCA protocol (37): briefly, samples were sonicated for 5 min in acetone, ethanol, methanol, and water, then immersed in a solution of H_2_O_2_ (30%) / NH_4_OH (30%) / H_2_O (1:1:5 by volume) and sonicated for 5 min at room temperature before soaking them for another 30 min at 60 °C. Afterwards, glass substrates were intensively rinsed with water, dried at 70 °C, and stored in a vacuum chamber.

Sample chambers were individually assembled by bonding the cleaned glass slides (25 x 75 mm, Menzel GmbH, Braunschweig, Germany) to plastic fluidic channels (μ-slide VI, ibidi, Munich, Germany). Polydimethylsiloxane produced from base and curing agent (SYLGARD184, Dow Corning Co., USA) served as bonding agent. Supported membranes were prepared by drying stock solutions of either DOPC or DPPC containing 0.2, 0.5, 0.8, 1.0 and 1.5 mol % biotin-DOPE in chloroform in a nitrogen stream. Dried samples were kept in a vacuum chamber at 25 °C for 12 h, and were re-suspended in de-ionized water to a concentration of ∼ 1 mg/ml. Small unilamellar vesicles (SUVs) were prepared by pulsed sonication of the lipid solution using a tip sonicator (Misonix, New York, USA) for 30 min at 1.0 W. Subsequently, SUV suspensions were incubated on the cleaned, bonded sample chambers where they formed a lipid bilayer on the glass surface. After 45 min. of incubation, the chambers were intensively rinsed with de-ionized water to remove remaining SUVs. Subsequently, the supported membranes were incubated with 1 μg/ml neutravidin solution for 30 min and were thoroughly washed with de-ionized water. Next, the membranes were incubated with aqueous CD95L-T4-btn (2μg/ml) solution for 1h at room temperature. Finally, the samples were carefully washed with pre-warmed cell culture medium and kept at 37 °C prior to cell experiments.

### CD95L containing supported membrane on silica beads

SUVs prepared as described above were used to form supported membrane on porous silica microspheres (NUCLEOSIL^®^ standard C18 phases, nonpolar from Macherey-Nagel, Düren, Germany). Lipid vesicles were incubated with silica beads at 60°C for 2h under rotation. The lipid-coated beads were collected by centrifugation and washed with PBS. CD95L functionalization of the supported membrane on the beads was performed as described above. Here, the beads were kept on a shaker during each incubation step.

### Cell imaging

Cell death was measured by microscopy using transmission imaging and visual cell morphology identification. Images were taken on a Leica SP5 confocal microscope (Leica Microsystems, Mannheim, Germany) or on an OlympusCKX41 wide-field microscope equipped with an Olympus PEN Lite CCD camera (Olympus Europa, Hamburg, Germany). Cell death kinetics were quantified from images taken every 10 minutes and by manually marking the first rounding and shrinkage event due to cell death. Marks were then automatically segmented and counted in ImageJ.

To obtain statistically reliable data, we collected results from 3 independent measurements for each experimental condition. For each experiment, cells in a concentration of 10^6^/ml cells were injected into each channel of the 6 channel chambers. The total number of cells was counted from 5 – 8 phase contrast images. Time-lapse images were recorded on a fully automated Leica SP5 confocal microscope (Leica Microsystems, Mannheim, Germany) equipped with an Apo 63x NA 1.4oil immersion objective using a proprietary macro for auto-focusing and multi-position imaging.

The microscope stage was maintained at 37 °C and CO2 at 5%. Apoptotic cells were identified by monitoring irreversible cell blebbing and fragmentation of cell nuclei. The area of cell adhesion and CD95-GFP clusters were detected by morphological identification and intensity threshold. Images were analyzed using ImageJ/Fiji (NIH,USA) as well as routines written in Matlab (R2013b, The Math Works Inc, MA, USA) following cell segmentation scripts (38).

### Quantification of CD95 receptor levels presented on cell surface

For the comparison of adhesion/apoptosis behaviors between different cells, the average amount of CD95receptors expressed on each cell type was calculated by FACS analysis using QIFIKIT (Dako) according to manufacturer’s protocol. Briefly, a calibration curve was constructed using calibration beads followed by determination of CD95 molecules on the cell surface. For CD95 staining, αApo-1 (Apogenix, 0.01 μg/μl) was incubated for 20 minutes on ice, followed by incubation of the secondary antibody (1:30 goat-anti-mouse phycoerythrin-conjugated; Dianova) for 30 minutes. Flow cytometry analysis was performed on a FACS Calibur (Becton Dickinson) using Cell Quest Software. The surface area of PanD24 and GBM cells was calculated from the projected diameter of non-adherent cells obtained from phase contrast images.

### Sphere formation assay

Neurosphere assays were performed as previously described with slight modifications (39). In brief, tumorspheres were harvested 3 to 7 days after culturing by centrifugation and dissociated using Accutase (Thermofisher), thereafter cells were centrifuged and Accutase was replaced by normal Neurobasal medium. The cell suspension was counted and 2×10^5^ cells per well were cultured on a 6 well plate. The next day, the newly formed spheres were collected by centrifugation, resuspended in 100μl of Neurobasal medium and were incubated for 4 h with either the ligand coated membrane or the control DOPC membrane. Afterwards the cells were harvested, centrifuged and the cell pellet was shortly treated with Accutase to obtain a single cell suspension. For the sphere formation assay 500 cells/well were seeded in a 96 well plate. From each condition 10 wells were plated and analyzed. The number of tumorspheres per well was scored by microscopic examination 10 days after seeding.

### Animal experiments

All animal experiments were performed in accordance with the institutional guidelines of the German Cancer Research Center and were approved by the Regierungspräsidium Karlsruhe, Germany.

### Subcutaneous injection of tumor cells and i.v. injection of coated beads

SCID beige mice (8-10 weeks old, female) were subcutaneously transplanted with 10000 GBM cells in 100 μl matrigel. Tumor size was measured every week with a caliper and after establishment of tumors, mice were intravenously injected with 2×10^6^ DOPC or 1% beads through the tail vein. Two weeks later animals were sacrificed and tumors were retrieved after transcardial perfusion.

### Orthotopic injection of tumor cells and coated beads

SMA-560 tumor cells were transplanted into 8 to 10 week old, female VMDK mice (in house breeding). Therefore, 5000 tumor cells were re-suspended in 2 μl of Hanks buffered saline solution (HBSS) and loaded into a 10 μl nanofill syringe (WPI) and stereotactically injected into the striatum (2.5 mm lateral of the bregma to a depth of 3mm) of isofluorane-anaesthetized mice. Five days post-injection 2×10^6^ control beads (DOPC) or CD95L-coated beads (1% Beads) were inoculated at exactly the same position as the tumor cells before. On reaching the no-go criteria, which are declined in the animal permission, mice were sacrificed and brains were retrieved after transcardial perfusion.

### Perfusion

Mice were anesthetized by intraperitoneal injection of 800 μl perfusion solution. After the thoracic cavity was opened, the heart was exposed and transcardial perfusion was performed with 10 ml HBSS. For immunohistochemistry, the mice were additionally perfused with 10 ml of 4% PFA.

### Immunohistochemistry of the brain

For immunohistochemistry of the tumor bearing brains, mice were sacrificed as described above and brains were extracted and fixed in 4% PFA overnight. 100 μm coronal sections were prepared using a vibratome (Leica, Germany). Sections were stored in PBS with 0.05% sodium azide until staining. For the staining, sections were incubated in PBS with 0.3% Triton X-100 and 5% horse serum for one hour to block unspecific binding sites. For different analyses, sections were stained with anti-phospho ERK (1:200; CST), anti-phospho Ser 473 Akt (1:25; CST), anti-cleaved Caspase-3 (1:100; CST), anti-Ki67 (1:200; Novus) and anti-pH3 (1:1000; Millipore) antibody diluted in blocking solution for 72 hours at 4°C with shaking. After washing the sections 3 times for 5 min each with PBS, secondary antibody staining was performed using the appropriate Alexa-fluorophore antibodies diluted in blocking buffer for 2h at room temperature. Furthermore, Hoechst 33342 was added to the secondary antibody dilution to stain for nuclei. Sections were then washed three times for 5 min with PBS, and in a last step with PB (Phosphate buffer without salt) buffer. After mounting with Fluoromount-G, the sections were allowed to dry at room temperature for overnight and were stored in the dark at 4°C until imaging.

---

**Figure S1.**
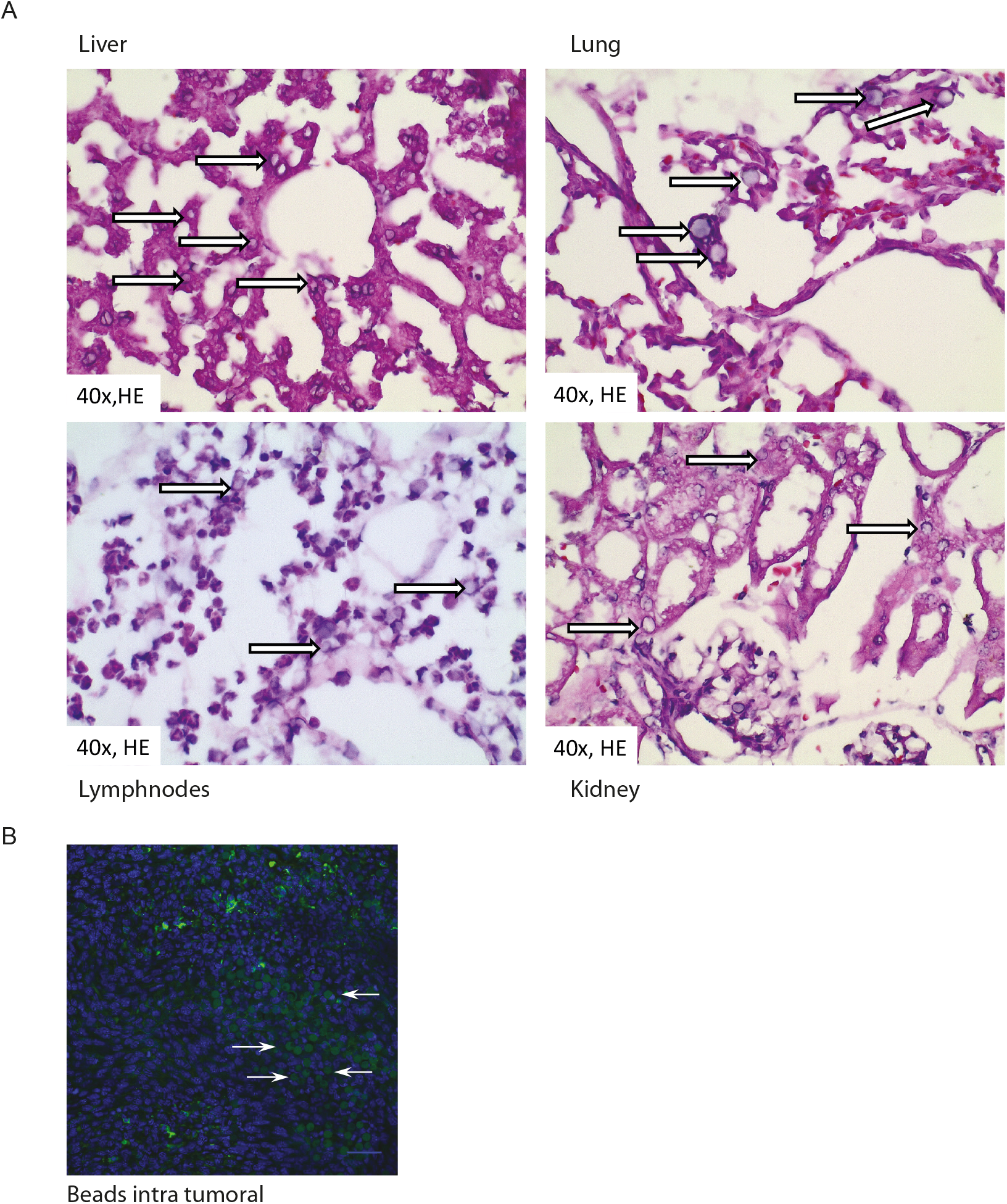
CD95L coated beads detection in organs and tumor. (A) The white arrows show the beads in H&E stained histological sections of liver, lung, lymphnode, kidney and (B) The white arrows show the beads in GBM tumors.

**Figure S2.**
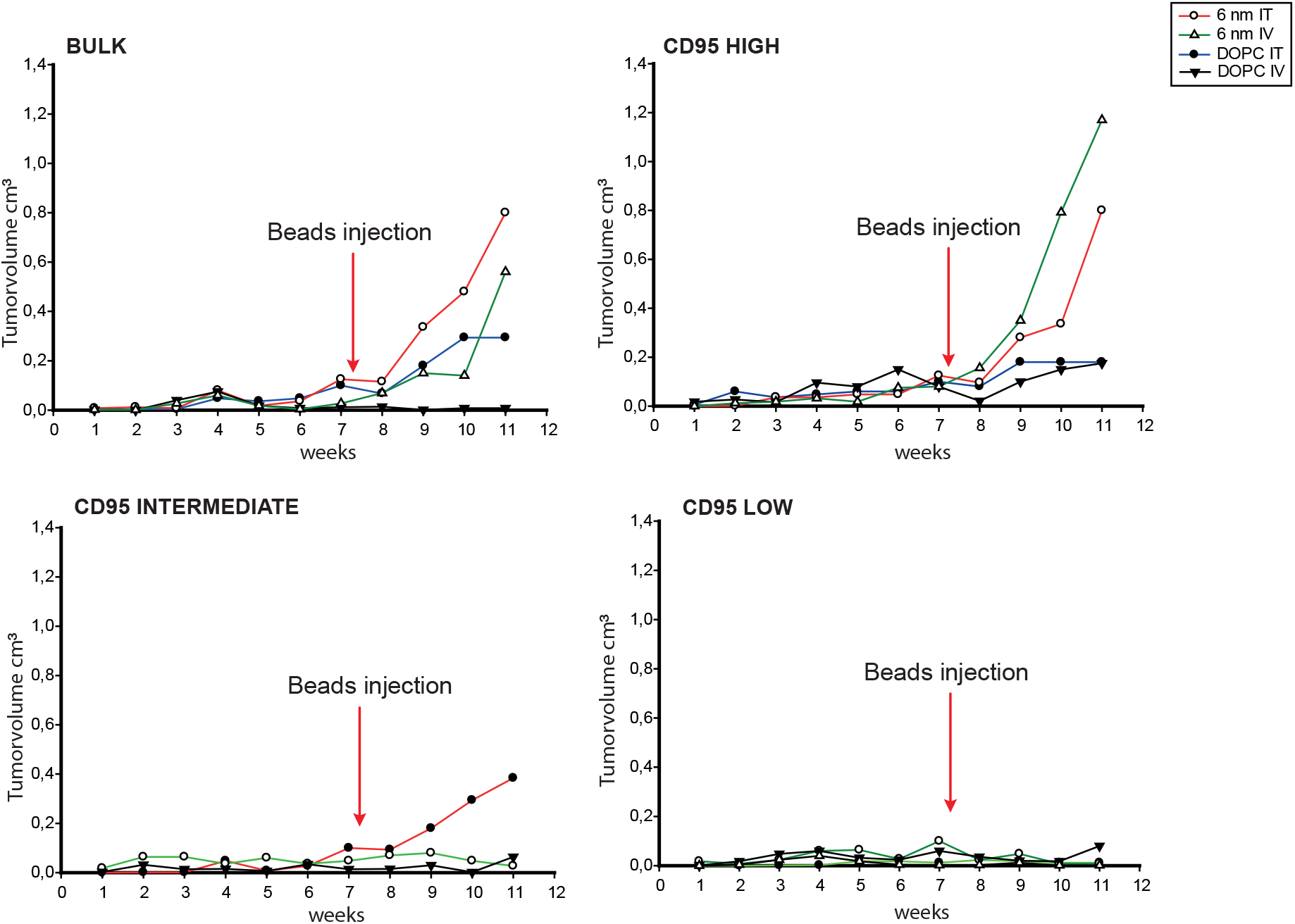
CD95L coated beads promote subcutaneous tumor growth. Tumor growth measured from 2^nd^ set of in vivo subcutaneous experiment from GBM10 cells.

**Figure S3.**
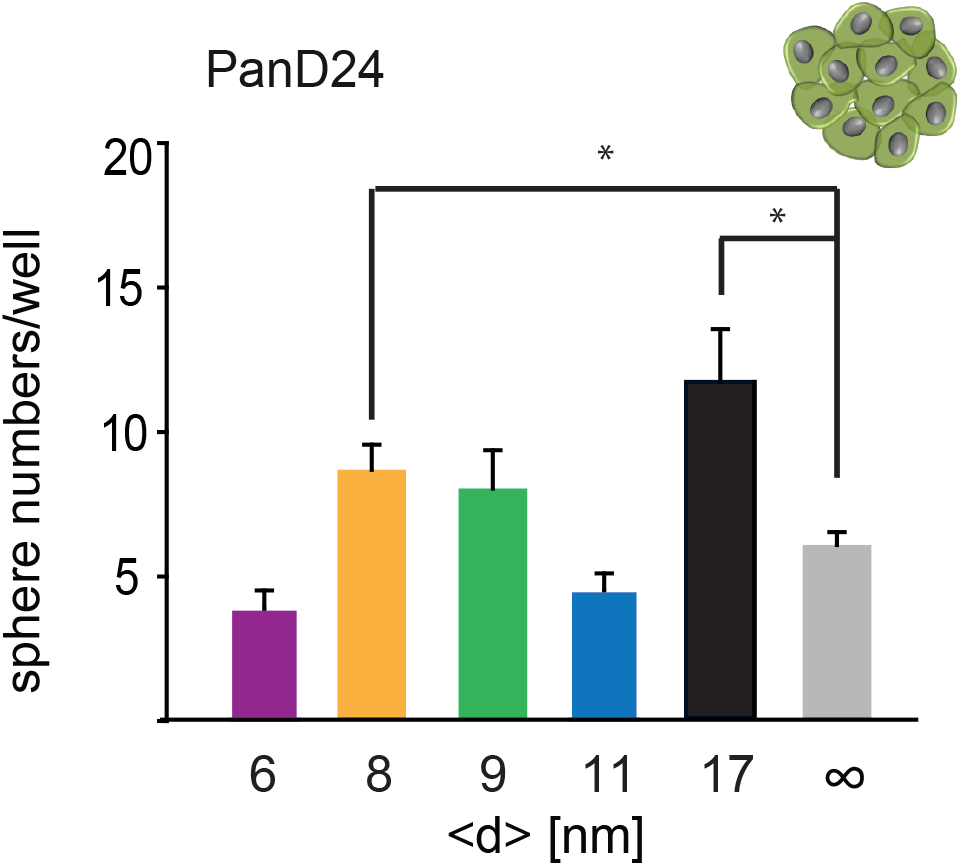
PDAC patient derived PanD24 sphere cultures were treated for 4 h with CD95L-coated membranes and were subsequently plated for sphere formation assessment. Values are mean ± S.D.; one-way ANOVA p* < 0.05.

**Figure S4.**
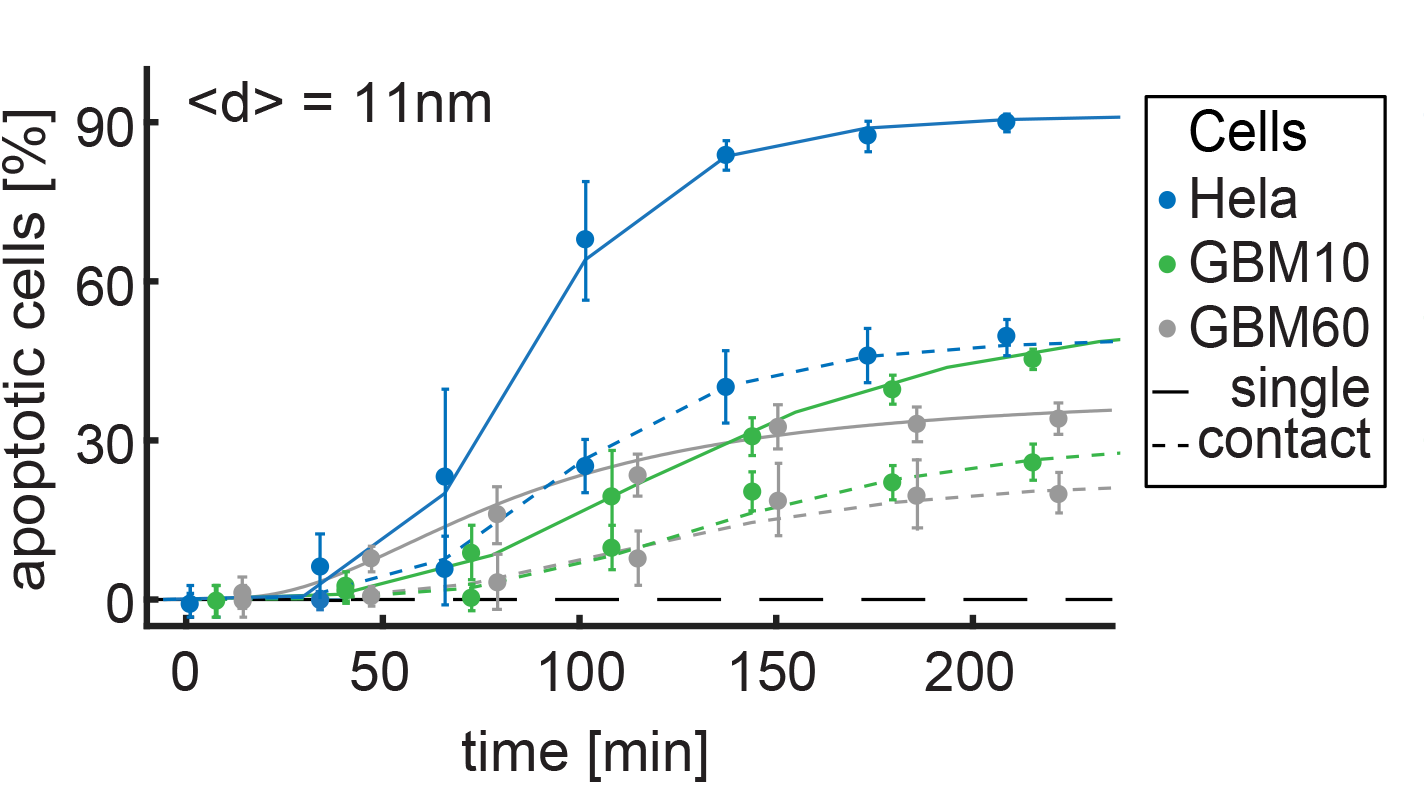
Cell-cell contact decreases the apoptotic activation compare to isolated cells. Cancer cells (specific color for each cancer cell line) incubated on CD95L membranes (<d>=11nm) and analyzed for the percentage of apoptotic cells according to having contact with other cells or not. Cells having contact with other cells (broken line) showed less apoptosis whereas isolated cells (solid line) showed more.

**Figure S5.**
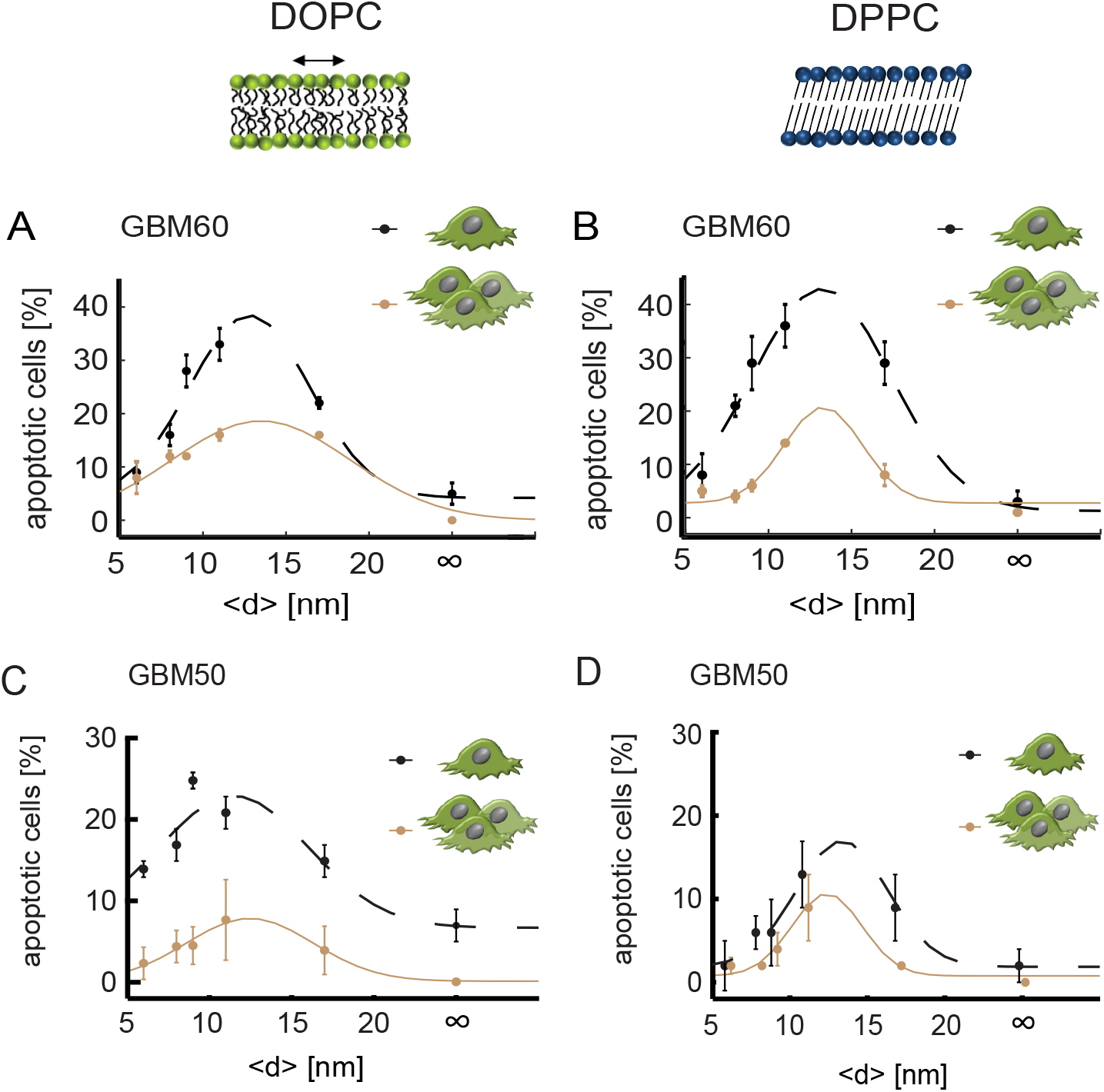
Cross-talk between cell-substrate and cell-cell contacts. At 37°C supported membranes composed of DOPC are in the fluid phase and enable CD95L mobility, whereas DPPC supported membranes are in the gel-like phase suppressing CD95L mobility. (A,C) Apoptotic behavior of GBM60 and GBM50 in contact with other cells on “fluid” DOPC membranes functionalized with CD95L at different <d> was analyzed at t = 200 min. A broken line indicates data of isolated individuals. n≥ 40 cells/condition. (B,D) Apoptotic behavior of GBM60 and GBM50 in contact with other cells on “solid” DPPC membranes (gel phase) functionalized with CD95L at different <d> was analyzed at t = 200 min. Broken line: data of isolated individuals. n≥ 40 cells/condition. Values are median ± M.A.D.; Significance levels according to p* * * < 0.001, p* * < 0.01, p* < 0.05.

**Table S1.**
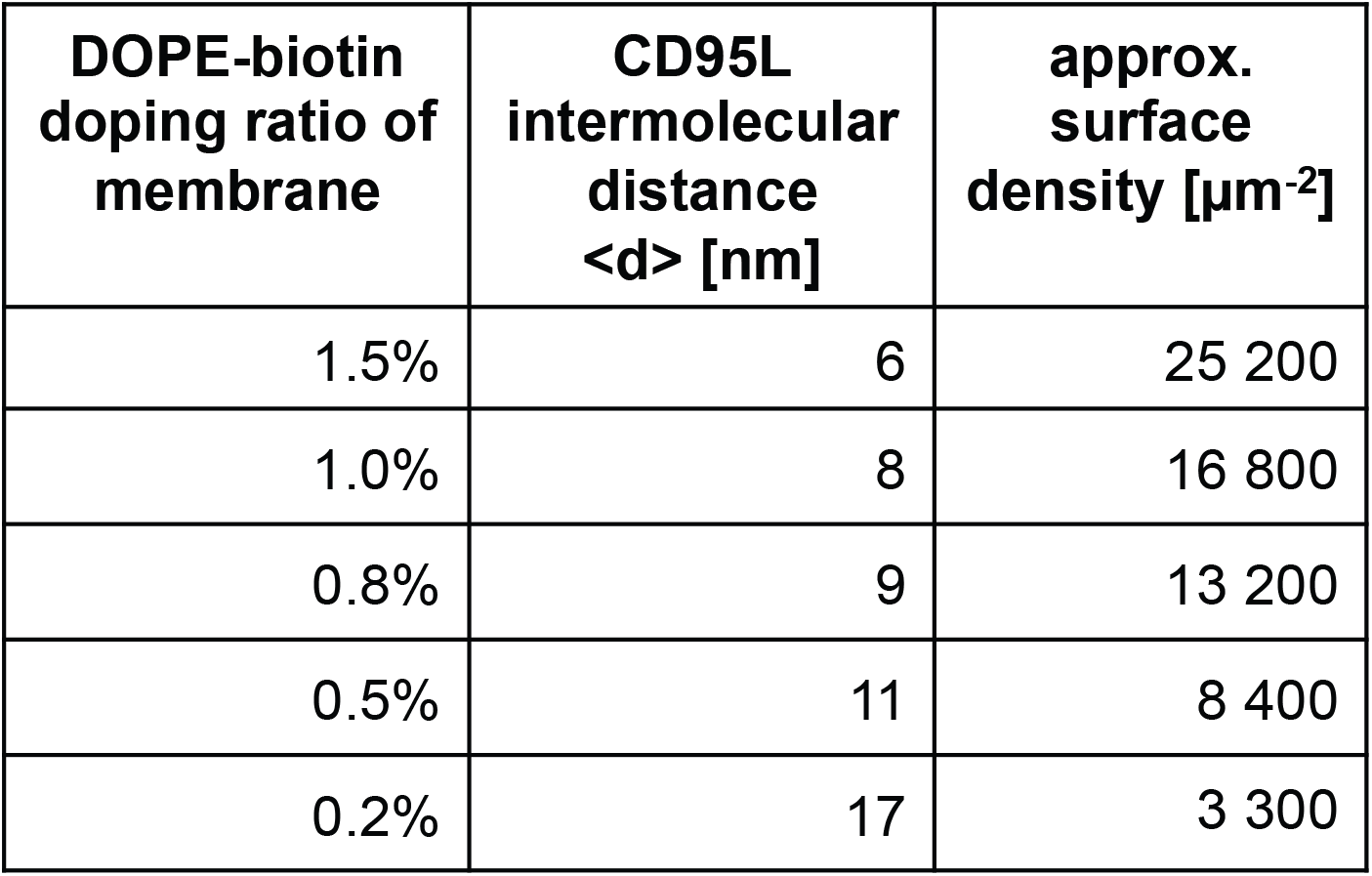
CD95L density on supported membranes.

**Table S2.** Result of Hill fits for PanD24 cells on DOPC membranes.

| **<d>**<br>**[nm]** | **max.**<br>**value [%]** | **half-time**<br>**[min]** | **n** |
| --- | --- | --- | --- |
| 6 | 66 | 25 | 1.8 |
| 8 | 77 | 39 | 2.1 |
| 9 | 92 | 36 | 1.5 |
| 11 | 72 | 34 | 1.8 |
| 17 | 46 | 45 | 2.0 |
| ∞ | 7 | 33 | 2.6 |

**Table S3.** Number of CD95 receptor molecules on different cancer cells.

| **Cell lines** | **# CD95/cell** | **approx. surface density [$\mu\text{m}^{-2}$]** |
| --- | --- | --- |
| PanD24 | 43 000 | 13.4 |
| GBM10 | 6 500 | 2.0 |
| GBM60 | 5 900 | 1.8 |
| GBM50 | 5 700 | 1.8 |
| Hela | 23 000 | 7.1 |

**Table S4.** Result of Hill fits for GBM cells on DOPC membranes.

| <d><br>[nm] | max.<br>value [%] | half-time<br>[min] | n |
| --- | --- | --- | --- |
| 6 | 14 | 100 | 2.4 |
| 8 | 21 | 44 | 1.8 |
| 9 | 27 | 65 | 1.7 |
| 11 | 21 | 46 | 1.8 |
| 17 | 17 | 92 | 2.6 |
| ∞ | 7 | 124 | 4.6 |

GBM60 on DOPC
| <d><br>[nm] | max.<br>value [%] | half-time<br>[min] | n |
| --- | --- | --- | --- |
| 6 | 9 | 80 | 3.6 |
| 8 | 20 | 81 | 2.5 |
| 9 | 20 | 72 | 2.1 |
| 11 | 29 | 68 | 1.7 |
| 17 | 22 | 80 | 2.9 |
| ∞ | 5 | 46 | 2.2 |

**Table S5.** Result of Hill fits for GBM cells on DPPC membranes.

| **<d>**<br>**[nm]** | **max.**<br>**value [%]** | **half-time**<br>**[min]** | **n** |
| --- | --- | --- | --- |
| 6 | 9 | 274 | 4.3 |
| 8 | 21 | 286 | 5.7 |
| 9 | 19 | 257 | 3.2 |
| 11 | 26 | 209 | 3.3 |
| 17 | 23 | 253 | 4.3 |
| ∞ | 7 | 235 | 5.7 |

**GBM60 on DPPC**
| **<d>**<br>**[nm]** | **max.**<br>**value [%]** | **half-time**<br>**[min]** | **n** |
| --- | --- | --- | --- |
| 6 | 11 | 177 | 4.3 |
| 8 | 37 | 184 | 3.4 |
| 9 | 35 | 123 | 3.7 |
| 11 | 47 | 245 | 3.2 |
| 17 | 41 | 158 | 2.5 |
| ∞ | 5 | 124 | 2.0 |

**Movie S1**. CD95 cluster formation over time on CD95L supported membranes (PanD24 cells expressing CD95-YFP).

**Movie S2**. GBM cells showed apoptosis as single whereas they survived as clusters upon CD95L supported membrane activation.

## References and Notes

1. M. R. Alderson et al., Fas ligand mediates activation-induced cell death in human T lymphocytes. J. Exp. Med. 181, 71–7 (1995).

2. J. Dhein, H. Walczak, C. Bäumler, K.-M. Debatin, P. H. Krammer, Autocrine T-cell suicide mediated by APO-1/(Fas/CD95). Nature. 373, 438–441 (1995).

3. T. Brunner et al., Cell-autonomous Fas (CD95)/Fas-ligand interaction mediates activation-induced apoptosis in T-cell hybridomas. Nature. 373, 441–444 (1995).

4. S.-T. Ju et al., Fas(CD95)/FasL interactions required for programmed cell death after T-cell activation. Nature. 373, 444–448 (1995).

5. A. Algeciras-Schimnich et al., Two CD95 tumor classes with different sensitivities to antitumor drugs. Proc. Natl. Acad. Sci. U. S. A. 100, 1144511450 (2003).

6. B. C. Barnhart et al., CD95 ligand induces motility and invasiveness of apoptosis-resistant tumor cells. EMBO J. 23, 3175–3185 (2004).

7. F. L. Scott et al., The Fas-FADD death domain complex structure unravels signalling by receptor clustering. Nature. 457, 1019–22 (2009).

8. K. E. Schlottmann, E. Gulbins, S. M. Lau, K. M. Coggeshall, Activation of Src-family tyrosine kinases during Fas-induced apoptosis. J. Leukoc. Biol. 60, 546–54 (1996).

9. A. Martin-Villalba, E. Llorens-Bobadilla, D. Wollny, CD95 in cancer: tool or target? TrendsMol. Med. 19, 329–335 (2013).

10. M. E. Peter et al., The role of CD95 and CD95 ligand in cancer. Cell Death Differ. 22, 549–59 (2015).

11. P. Legembre, B. C. Barnhart, M. E. Peter, The relevance of NF-kappaB for CD95 signaling in tumor cells. Cell Cycle. 3, 1235–9 (2004).

12. S. Kleber et al., Yes and PI3K bind CD95 to signal invasion of glioblastoma. Cancer Cell. 13, 235–48 (2008).

13. I. Sancho-Martinez, A. Martin-Villalba, Tyrosine phosphorylation and CD95: A FAScinating switch. Cell Cycle. 8, 838–842 (2009).

14. M. Tanaka, T. Suda, T. Takahashi, S. Nagata, Expression of the functional soluble form of human fas ligand in activated lymphocytes. EMBO J. 14, 1129–35 (1995).

15. T. Suda, H. Hashimoto, M. Tanaka, T. Ochi, S. Nagata, Membrane Fas ligand kills human peripheral blood T lymphocytes, and soluble Fas ligand blocks the killing. J. Exp. Med. 186, 2045–50 (1997).

16. P. Schneider et al., Conversion of membrane-bound Fas(CD95) ligand to its soluble form is associated with downregulation of its proapoptotic activity and loss of liver toxicity. J. Exp. Med. 187, 1205–13 (1998).

17. L. A. O’ Reilly et al., Membrane-bound Fas ligand only is essential for Fas-induced apoptosis. Nature. 461, 659–63 (2009).

18. M. Tanaka, E. Sackmann, Polymer-supported membranes as models of the cell surface. Nature. 437 (2005), pp. 656–663.

19. A. S. Burk et al., Quantifying Adhesion Mechanisms and Dynamics of Human Hematopoietic Stem and Progenitor Cells. Sci. Rep. 5, 9370 (2015).

20. M. Drachsler et al., CD95 maintains stem cell-like and non-classical EMT programs in primary human glioblastoma cells. Cell Death Dis. 7, e2209 (2016).

21. R. B. Gennis, Biomembranes: molecular structure and function (Springer-Verlag, New York, 1989).

22. L. Gao et al., Endothelial cell-derived CD95 ligand serves as a chemokine in induction of neutrophil slow rolling and adhesion. Elife. 5, e18542 (2016).

23. M. Teodorczyk et al., CD95 promotes metastatic spread via Sck in pancreatic ductal adenocarcinoma. Cell Death Differ. 22, 1192–1202 (2015).

24. A. C. Cruz et al., Fas/CD95 prevents autoimmunity independently of lipid raft localization and efficient apoptosis induction. Nat. Commun. 7, 13895 (2016).

25. J. Chen, S. C. Almo, Y. Wu, General principles of binding between cell surface receptors and multi-specific ligands: A computational study. PLOS Comput. Biol. 13, e1005805 (2017).

26. C. M. Nelson, C. S. Chen, Cell-cell signaling by direct contact increases cell proliferation via a PI3K-dependent signal. FEBSLett. 514, 238–242 (2002).

27. R. Levayer, B. Hauert, E. Moreno, Cell mixing induced by myc is required for competitive tissue invasion and destruction. Nature. 524, 476–480 (2015).

28. W. Stallaert, O. Sabet, Y. Bruggemann, L. Baak, P. Bastiaens, Contact inhibitory Eph signaling suppresses EGF-promoted cell migration by decoupling EGFR activity from vesicular recycling. bioRxiv, 202705 (2018).

29. A. Grakoui et al., The immunological synapse: a molecular machine controlling T cell activation. Science. 285, 221–227 (1999).

30. M. Barthélémy, S. V. Buldyrev, S. Havlin, H. E. Stanley, Multifractal properties of the random resistor network. Phys. Rev. E. 61, R3283–R3286 (2000).

31. J. R. James, R. D. Vale, Biophysical Mechanism of T Cell Receptor Triggering. Nature. 487, 64–69 (2012).

32. F. Aoudjit, K. Vuori, Engagement of the α2β1 integrin inhibits Fas ligand expression and activation-induced cell death in T cells in a focal adhesion kinase-dependent manner. 95, 2044–2052 (2016).

33. T. G. Bivona et al., FAS and NF-κB signalling modulate dependence of lung cancers on mutant EGFR. Nature. 471, 523–6 (2011).

34. W. Wick et al., A phase II, randomized, study of weekly APG101+reirradiation versus reirradiation in progressive glioblastoma. Clin. Cancer Res. 20, 6304–13 (2014).

35. A. M. Xu, P. H. Huang, Receptor tyrosine kinase coactivation networks in cancer. Cancer Res. 70, 3857–60 (2010).

36. C. Zuliani et al., Control of neuronal branching by the death receptor CD95 (Fas/Apo-1). Cell Death Differ. 13, 31–40 (2006).

37. W. Kern, D. Puotinen, Cleaning solutions based on hydrogen peroxide for use in silicon semiconductor technology. RCA Rev. 31, 187–206 (1983).

38. Cornelia Monzel and Christoph Möhl, in Bioimage Data Analysis, Kota Miura, Ed. (Wiley-VCH, Weinheim, 2016), pp. 63–97.

39. S. R. Ferrón et al., A combined ex/in vivo assay to detect effects of exogenously added factors in neural stem cells. Nat. Protoc. 2, 849–859 (2007).

